# Investigation of Different Free Image Analysis Software for High-Throughput Droplet Detection

**DOI:** 10.1101/2021.04.13.439618

**Authors:** Immanuel Sanka, Simona Bartkova, Pille Pata, Olli-Pekka Smolander, Ott Scheler

## Abstract

Droplet microfluidics has revealed innovative strategies in biology and chemistry. This advancement has delivered novel quantification methods, such as digital droplet polymerase chain reaction (ddPCR) and antibiotic heteroresistance analysis tool. For droplet analysis, researchers often use imaging techniques. Unfortunately, the analysis of images may require specific tools or programming skills to produce the expected results. In order to address the issue, we explore the potential use of standalone freely available software to detect droplets. We select four most popular software and classify them into instinctive and objective types based on their operation logic. We test and evaluate the software’s i) ability to detect droplets, ii) accuracy and precision, and iii) overall user-friendliness. In our experimental setting we find the objective type of software is better suited for droplet detection. The objective type of software also has simpler workflow or pipeline, especially aimed for non-experienced user. In our case, CellProfiler^™^ (CP) offers the most user-friendly experience for both single image and batch processing analysis.

---

Droplet microfluidics has become a powerful tool for high-throughput analysis over the last decades^1^. Compartmentalization of samples and massive parallelization of experiments suitable for excellent statistical data, screening of large numbers of compounds, or for seeking rare events in large pools of molecules or organisms^2^. Droplets are often applied for high sensitivity nucleic acid diagnostics^3^ or microbiological studies^4^.

[Imaging is a popular method to analyze droplet microfluidic experiments]. This method helps researchers in practical detection, such as bacterial surveillance of foodborne contamination^5^, screening specific substrates^6^, single cell analysis^7^ and detecting viable bacteria utilizing smartphone^8^. Over four thousand articles have been published during the last decade related to imaging and droplet microfluidics^9^. This shows that imaging is a popular technique, either with brightfield or fluorescence microscopy^10^. Though, imaging techniques require image analysis tools to extract and interpret the results.

[Image analysis (IA) tools are available for different scenarios, from single image to real-time counting analysis]. These diverse tools have been used for many different analyses, such as detecting droplets^11^ and screening protein crystals within droplets^12^. Moreover, droplet microfluidics have been developed as frontier techniques in chemistry and biology, e.g. for absolute DNA quantification by Digital Droplet Polymerase Chain Reaction (ddPCR)^13,14^ or detecting viable bacteria and hetero-resistance in antimicrobial experiment^15,16^.

[However, implementing IA for droplet analysis often requires specific skills in programming and the methods are not widely available in non-specialist laboratories]. Most of the published articles in droplet detection use scripted programs, such as Circular Hough Transform in Python programming language^17^, Mathematica^18,19^, Scikit-image in Python^20^, Image Processing Toolbox from MATLAB^21^, and OpenCV in C++ ^22^. There are some user-friendly software and may be used for droplet microfluidics image analysis, such as Zen imaging program^23^ and NIS-Elements from NIKON^5^. Yet, these kinds of programs are only commercially available.

[There is a need for widely accessible and user-friendly IA tools for droplet analysis]. Open-source software is available and can be used to detect and/or analyze droplets. Such software are for example FIJI/ImageJ which has been used to analyze image data in general^24^ or droplets^25^, or CellProfiler which was developed to identify and measure various bioimage data^26^. Even though some published articles mention the use of the software, there are limited information regarding their workflow (the data is often missing from publications). Moreover, novel workflows can be constructed by combining functions, modules or pipelines from different software, like a puzzle^27^. [Here, we i) demonstrate how to use different software for the analysis of droplet images and ii) explore the differences and similarities of workflow logic in the different software].

## Results and Discussion

[Most popular software for image analysis are ImageJ, CP, Ilastik and QuPath]. To find the popularity, we executed Twint^28^ Python script using each of the software’s name as the keywords and find through hashtags (#). The keywords were also used for finding the popularity in published articles in Scopus’ repository. Our search showed that most popular software are ImageJ^24^, CellProfiler^29^, Ilastik and QuPath (Figure 1a). Ilastik is a machine learning based image analysis software^30^ and QuPath has been used as whole slide image analysis tool^32^. We continued with these four popular software tools and used them to detect droplets on images previously described by Bartkova et al.^31^ (Figure 1b). Then, we took a deeper look into the working logic of the software and assessed their performance with different key parameters (Figure 1c).

**Figure 1.**
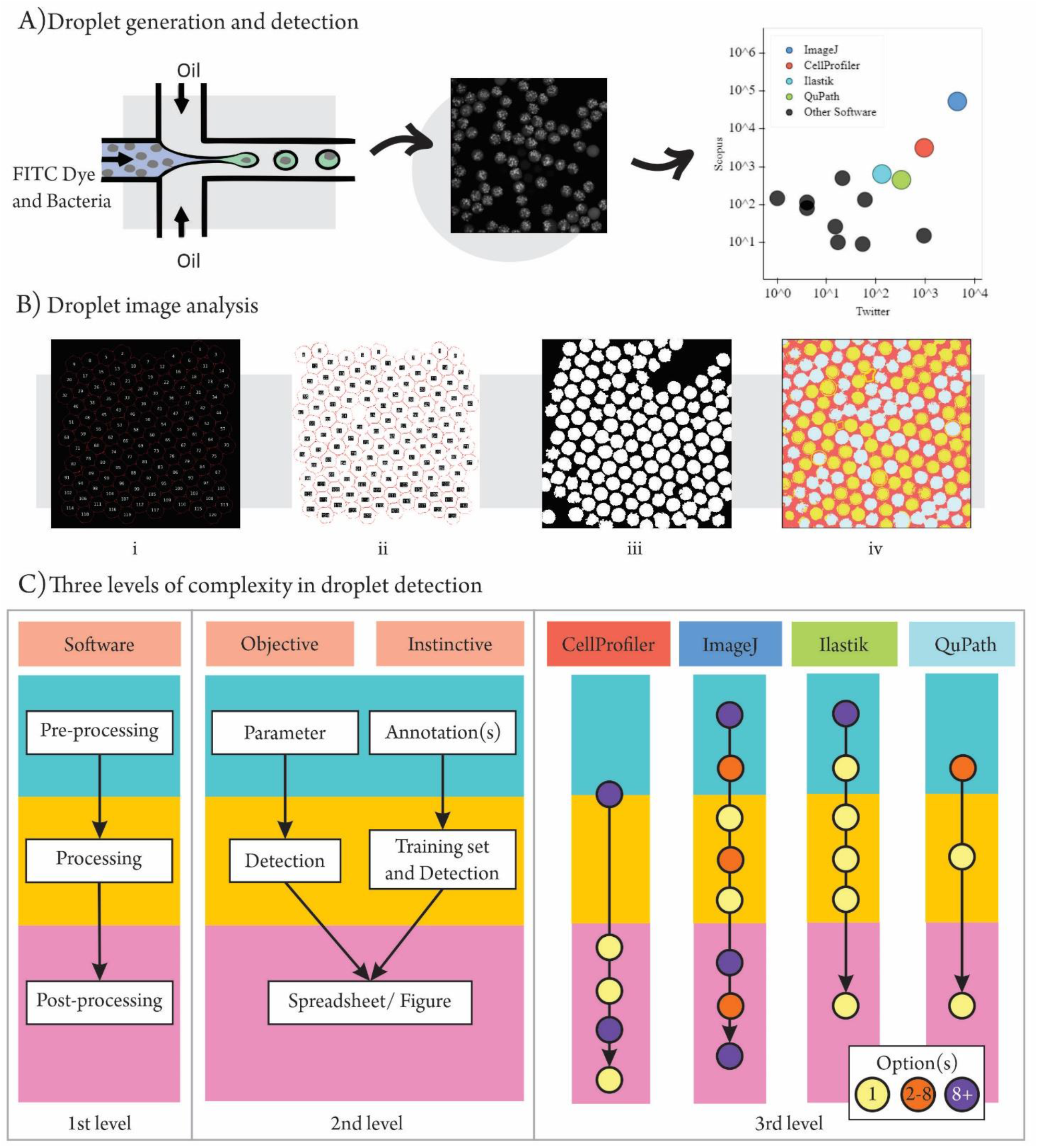
Schematic of droplet generation and image analysis. (A) We generated water-in-oil droplets using flow-focusing microfluidic chip as described previously by Bartkova et al.^31^ (left). We used fluorescent microscope to obtain “Raw images” of droplets that contained fluorescence producing bacteria (middle). For the analysis of droplet images we used four most popular image analysis that were selected according to hits in social media (Twitter) and Scopus Search (obtained on February 11th, 2021) (right). (B) Droplet detection comparison between: i) ImageJ, ii) Cellprofiler, iii) Ilastik and iv) QuPath. (C) We divide the image processing software into two groups (Objective and Instinctive) and explore their logic and working principle on three levels of complexity. 1) The first level shows that used software are very similar in their basic image processing logic. They usually have three processing stages in their image analysis logic: pre-processing, processing, and post-processing. 2) The second level shows distinction between two groups of software in droplet detection: Objective, where user defines how to detect droplets by giving specific parameters (e.g., threshold or size) or Instinctive, where user classifies/annotates grouping of pixels on image. 3) The third level shows number of different steps and modules in processing stages. Purple circle = more than 8 options are available, orange circle = 2-8 options, and yellow circle = only 1 option is available.

[We divided the selected software into two groups (objective and instinctive) according to their logic and principle]. In the objective software group (CP and ImageJ) user has to manually provide settings for the program to select the pixels of interest with numeric or known parameter in order to detect droplets. In the instinctive group (ilastik and QuPath), on the other hand, the user may select the areas of image and manually annotate them as objects of interest (e.g. droplets or background) for pixel classification. Based on these characteristics, we described the complexity of the process with three increasing levels and used it to direct the droplet detection.

We used terms i) pre-processing, ii) processing and iii) postprocessing. i) In pre-processing, we modified, adjusted, and prepared the image data for further use. For instance, we performed pre-processing to duplicate the image data or introduce features on the image.

ii) In processing, we conducted segmentation or pixel partitioning based on color, intensity or texture along with droplet detection or counting process^33^. Usually, processing steps may help the user to obtain specific type of data^34^. In our case, we introduced thresholding to distinguish between the background (dark) and the foreground (droplets). For the details, CellProfiler came handy and only needed one module named “IdentifyPrimaryObject” which contained some options to detect droplets. This included thresholding, smoothing, segmentation, and automatic selection. In ImageJ, processing steps had three options: “Thresholding”, “Watershed” and “Analyze particle”. Similar to CP, these three steps will provide selections to detect the droplets. In the processing part, Ilastik had to process “Thresholding”, “Object Feature Selection”, and “Object Classification” for selecting the droplets and discarded the background. In QuPath, we found all of these features in “Pixel Classifier”. The settings included classifier from Artificial Neural Network with Multi-Layer Perception (ANN_MLP)^35^ with high resolution, using four multiscale features (Gaussian, Gradient Magnitude, Hessian determinant, and Hessian Max Eigenvalue) with probability as an output.

iii) For the last step, in post-processing, we prepared data extraction or generation for further use, for example to generate table of data or type of images for visualization. In CellProfiler, this last step was performed with “OverlayOutlines”, “OverlayObject”, “Dis-playDataOnlmage”, “ExportToSpreadsheet”. These modules generated the images and results in. CSV format. The order was similar in ImageJ and Ilastik but the option was available in “ROI Manager” and “Export”, respectively. In QuPath, the results can be obtained by exporting annotations from detected objects or called as labeled image. We used Groovy script to generate this result using commands in “Workflow” tab. For brief workflow/ pipeline, we provide the scheme of third level complexity in Supp. Fig. SI.

[CP has more than 96% accuracy in detecting droplets]. By comparing the results with golden truth (7145 droplets by manual count), we investigated the ability of analyzed software to detect droplets. We only counted droplets that did not touch the image border and did not make a bundle (joint droplets because of failed segmentation). We performed sensitivity and specificity test using True Positive (TP), False Positive (FP) and False Negative (FN) values based on the comparison with manual counting^36^. The TP confirms the positive droplet detection in the data. For FP, the value is obtained by finding false droplet detection or underestimation (type I error). In FN, the software does not detect the droplet or performs overestimation (type II error). We defined TN as background (black = 0). After the calculation, we obtained the accuracy ((TP+TN)/(TP+TN+FP+FN)) and precision (TP/(TN+TP)) from the detection. This accuracy explains the ratio between the correct droplet detection and total number of droplet detection. On the other hand, precision describes the probability to produce the correct droplet detection in total positive detection^37–39^. The accuracy of each detection ranging from 74.7% up to 96.2%. One of the software managed to generate precision up to 99.8% (Supp. Table 1).

From the Figure 2, we can see how each group share similar errors in every event (detection per image). We compared the false detection results (both FP and FN) from each of the software. We found that the objective group (CP and ImageJ) have less false detection compared to the instinctive group (ilastik and QuPath). However, Ilastik and QuPath received high error because they do not have filter to eliminate the droplets which touch the border, and some droplets are falsely detected as joint droplets (Supp. Fig. 2). The Figure 2 also indicates the image which may have bad quality for droplet detection. For instance, the image number 2, 19 and 64 depict the highest error values from all four software. Notwithstanding, CP outperforms the other software and has both high accuracy and precision.

**Figure 2.**
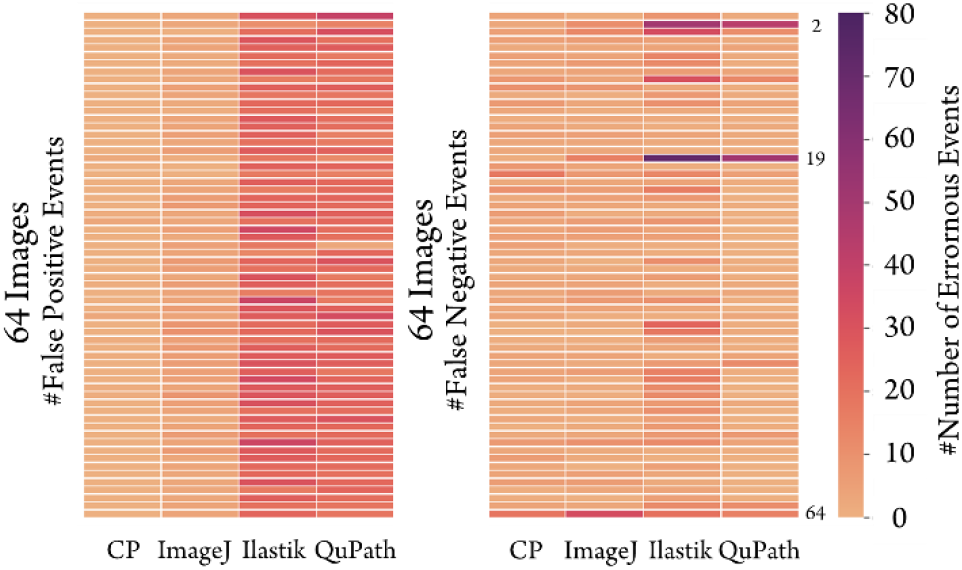
Droplet detection errors are higher in instinctive software in the diagnostic test. The figure shows the False Positive (FP, wrong detected droplet) and False Negative (FN, wrong undetected droplet) events per image. Each block represents the event or image error detection. The scale shows the number of errors (dark = high, bright = low).

[CP is the most suitable software for batch analysis or high throughput analysis]. In CP, we can analyze a whole set of images with a press of one single button “Analyze Images” on the main menu. The software will process available images uploaded in the “Images” module (default module). We tested and used the batch processing option to analyze 64 images straight after we had our pipeline/workflow set. In ImageJ, we processed the batch analysis using recorded macro from single image analysis. We also performed some macro script cleaning (e.g., closing unnecessary tabs during the process) which was written in the macro recorder. After the cleaning, we selected the input and output folder and performed the batch processing through the “Process” tab. For Ilastik, we executed the batch processing after the last option of the pipeline. We just needed to upload the images and started the “Process all files”. QuPath demanded macro programming commands for executing batch analysis. However, this software provided automated script generator which simplified the macro record to perform batch analysis. ImageJ and QuPath required macro script for batch analysis. Even though this macro script was easy to do, creating macro script for the first time could become an obstacle for researchers who are not familiar with any programming language or practices^40^. From our viewpoint, CP and Ilastik had user-friendly interface for batch processing and did not require any programming steps.

[Objective class software are flexible and have modular options to process image(s)]. As objective tools, CP and ImageJ offered options which could be added and removed depending on the user’s preferences, such as type of thresholding algorithm, filters, and other modules. In instinctive software, the features were embedded in the pipeline and had limited availability for additional settings. For example, Ilastik had some pre-defined pipelines: one of them was object classification and pixels classification^30^. These two were fixed in the interface of Ilastik and might be re-arranged only through Python programming. From the Figure 1(C), the third level of complexity also represents the modularity in which Ilastik and QuPath were more limited than CP and ImageJ. For instance, CP had “IdentifyPrimaryObject” module which could be duplicated in one pipeline, while in Ilastik, “Thresholding” could be performed only once within the pre-defined workflow. This complication placed Ilastik as the least flexible tool, followed by QuPath.

Macro programming language affects the software processing time, particularly in batch analysis]. CP or ImageJ expected less memory for use since they did not implement machine learning classification methods in our pipelines. The objective software used object logic classification^41^ and did not require training set to test the defined parameter, e.g. size of the object or maximum length of the object. In QuPath and Ilastik, the classification depended on a supervised machine learning process^30,41^. We used the manual annotations (droplets and background) to make the classifier before processing the whole pixels. We also compared the minimum hardware requirements for each of the tools (Supp. Table 2). Based on the comparison, ImageJ was the only one which did not put any minimum requirement for the Random-Access Memory (RAM). We also expected that the instinctive software might take more time to process the whole set of images. Therefore, we also tried to run the whole pipeline and compared the performance from each of the tools. We tested each pipeline with the same computer having Intel^®^ Core^™^ i3-9100F processor, 8Gb RAM, NVDIA GeForce GTX 1660 SUPER and 120Gb SSD PANTHER We found that CP and Ilastik had running time 978s and 765s while ImageJ and QuPath only needed 98s and 38s. Tool’s batch processing language (macros) may cause this difference. At the beginning, we expected Ilastik and QuPath to have longer processing time than CP and ImageJ because of the machine learning based processing. Yet, ImageJ and QuPath performed faster than others. In principle, there are two types of program which bioinformaticians use: compiled and interpreted^42^. ImageJ and QuPath use Java based (macros) code which is compiled once before the program process the batch analysis. This allows the program to run faster. On the other hand, CellProfiler and Ilastik use Python to process batch analysis. In Python, variables and functions will be run through interpreter every time the program needs to process the task, in our case to detect droplets in every image. Therefore, we presume that these differences in the implementation of batch processing might be the reason for these differences in processing times.

[CellProfiler and ImageJ have sufficient examples and documentation for novice user]. Each of the software provide documentation and examples for guiding their user. CellProfiler and ImageJ have been developed since 1987 and 2005, respectively^40,43^. Therefore, these objective software have more users and examples, e.g. ImageJ has a distribution for compiling the biological image analysis plugins called Fiji^24^. CP also provides some tutorials, examples, and other documentations on their website, e.g. detecting different cells morphology and tracking objects (www.cellprofiler.org). On contrary, Ilastik and QuPath have limited documentation for accompanying new users. However, there are some forums such as im-age.sc forum (forum.image.sc) which are actively helping other bioimage researchers or software users.

[Plugins in CellProfiler and ImageJ can be used as an extensible option in processing image]. Plugins or add-on can be used to improve default options within the software. This may be utilized by other software developers. As an additional option, plugins may help the user to implement specific cases of detection. Before Ilastik and QuPath were developed, ImageJ had plugins called Trainable WEKA Segmentation which in principle works similarly to instinctive software^44^. In CellProfiler, plugins are also available. For instance, we found one plugin to analyze Mass Cytometry (multiplexed images), called ImcPluginsCP^45^. Here, we did not add any plugins to detect droplets and we used similar settings to see the tool’s ability to detect and count the droplets. The extension software for CellProfiler, CellProfiler Analyst (CPA)^46^, could be an option to enhance droplet detection and have been described briefly our previous research^31^. Based on our classification, CPA belongs to instinctive software because users need to supervise or train the data at the beginning. However, this software is not a standalone software and requires feature extraction or properties file which contain the observed data from CP^31,47^.

[Objective software are more suitable for analyzing droplet microfluidics image data]. The objective software provide more options e.g. to disregard the object which touches the border/frame which resulted on high accuracy and precision. On the other hand, the instinctive software required more optimization to train the classifier. We only used 12 lines (5 lines for determining droplets and 7 lines to define borders between droplets and background) to supervise each class (background and droplet). Each line represents the pixels for each group. This pixel manual selection works better if the image has similar properties in majority and represent the pixels distribution of an object, for example, borders between droplets and the empty droplets. Even though the droplet’s border looks the same across the image, the pixel distributions are varied. We picked more lines to define borders. However, this cannot represent all of the properties and may result joint droplets. To overcome this, larger training set and improvement of classifier would, presumably, give better result. As an image processing tool, the instinctive software like QuPath has a more specific purpose. This software was created to accommodate whole slide image and large image data analysis, specifically for complex tissue images^32^. Yet, there has been made a comparison between QuPath and CellProfiler coupled with CellProfiler Analyst^41^. The comparison also shows the pro and cons between the objective and instinctive software in renal tissue. Furthermore, ImageJ, CellProfiler, Ilastik and QuPath have shown their capability in detecting droplets and generating the results as standalone tools. Based on user-friendliness, which is rated in the Table 1, CellProfiler is the easiest tool to use for droplet detection.

**Table 1.**
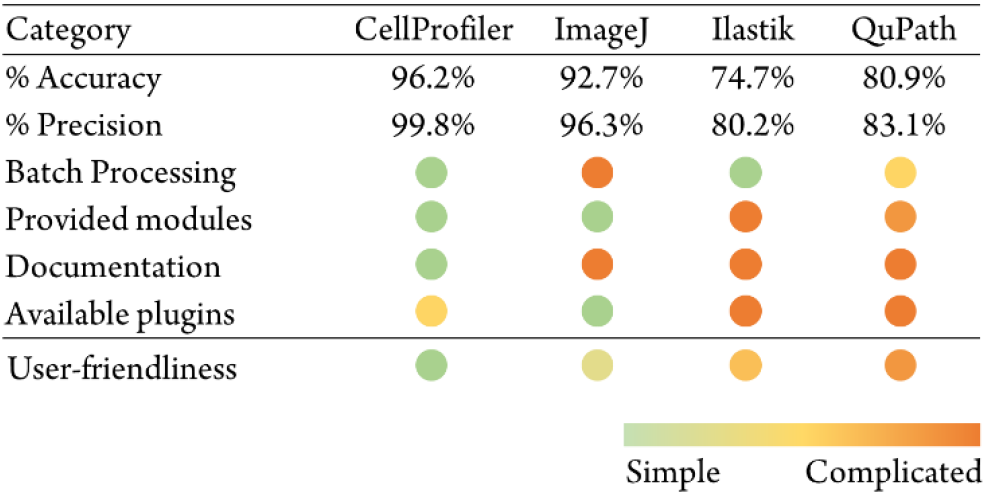
CP provides more simplicity among other software

Droplet detection is often used as preliminary step in droplet microfluidic experiment. It is possible to expand the pipeline for further analysis e.g. bacteria detection^31^, enzyme reaction measurement^48^, chemical purification analysis^49^, and metal extraction^50^. This step is usually performed to extract the different aspects of droplet (size, texture, volume and etc.) through pixel analysis. Yet, each software has their own option and feature to obtain the particular information, for example, “MeasureObjectSizeAndShape” and “MeasureObjectIntensity” in CP and “Set measurement” and “ROI Manager” in ImageJ. Nonetheless, this further analysis is not in the scope of this article. We try to focus on the principle of droplet detection in different software and assess the user-friendliness.

## Conclusion

This investigation gives insights to processing droplet microfluidic images using the four currently most popular software tools. We classified types of open-source software into objective and instinctive group. Both groups have three levels of complexity that cover pre-processing, processing, and post-processing steps. These steps help users, specifically with no programming experience, to choose and perform their image analysis. In our experimental setup we found that the objective type of software is better suited for droplet analysis. The objective type tools also have simpler workflow or pipeline, especially aimed for non-experienced user. Considering all of the aspects, in our case, CellProfiler^™^ offered the most user-friendly and accurate experience for droplet detection. However, the optimal software choice may definitely be different for other users depending on their experiment conditions and acquired images. Our paper would serve as a starting point for them to compare available solutions and start with settings optimization. In addition, published research, documentation, or forum discussions (such as www.image.sc) helps finding the most suitable software pipeline for droplet image analysis.

## Methods

### Software search and selection

We used selected software tools to detect droplets using procedure explained by Bartkova et al^31^. We found several available and accessible software tools online such as CellProfiler^29^, ImageJ^24^, Ilastik^30^, QuPath^32^, Icy^51^, BioFilmQf^52^, CellOrganizer©^53^, CellCognition^54^, BioImageXD^55^, BacStalk^56^, Advanced CellClassifier^57^, Phenoripper^58^, and Cytomine©^59^. To find the most preferred tools, we used Twint - Twitter Intelligence Tool script^28^ written in Python and used each tool’s name as a keyword. The processing was executed in Jupyter Notebook (ver. 6.0.3)^60^ within Anaconda Navigator^61^. We also imported datetime and pandas as additional libraries. The Scopus search was performed using same keyword and both of the results were visualized using bokeh and numpy library in Python^62 64^.

### Droplet generation and image acquisition

We repeated the method described in Bartkova et al. to generate droplets and their image data^31^. We used a set of 64 images to test the most popular software to detect the droplets. Using the data, we calculated accuracy by comparing the results with manual counting.

### Image analysis with most popular software

The image data were analyzed first as a single image using ImageJ (ver. l.52p), CellProfiler (ver. 4.0.3), Ilastik (vet. 1.3.3), and QuPath (ver. 0.2.3). We used our previous pipeline in CellProfiler as a basis to explore the other tools. We used the “IdentifyPrimaryObject” to detect droplets. We also used the same setting which also provided in our GitHub repository (github.com/taltechmicrofluidics/CP-for-droplet-analysis). The “MeasureObjectlntensity” and “ExportToSpreadSheet” modules were also set as previously. The results were obtained automatically after pressing “Analyze Image” button.

For ImageJ, we recorded the workflow in the macro record option. This record was used to make scripts for batch processing. The parameter was set within “Set Measurement” under “Analyze” tab and we only ticked “Area” for obtaining the pixels’ area in one droplet. This followed with processing workflow, which included segmentation using “Threshold” under “Adjust” option in “Image” tab. The threshold was determined as 1507, corresponding to 0.023 scale, described in our previous article using CP. The thresholding was followed with “Watershed” to separate droplets from each other. The counting was performed using “Analyze Particles” under “Analyze” tab. We set the size corresponding to the range we described in CellProfiler, 22500 up to 62500 pixel^2^ with 0 circularity. Once we finished the processing step, we downloaded the image through “Flatten” option in the “ROI Manager” menu. We obtained the results in the table which appeared straight after we performed the analysis.

In Ilastik, we used “Pixel Classification” and “Object Classification” pre-defined workflow. We loaded the image and selected the features for the training set. Since we did not have any reference regarding this type of workflow, we used the recommendation from image.sc forum, starting by adding 0.30, 1.00 and 3.50 sigma or scale correspond to the selected features e.g. Gaussian Filter, for color/intensity, edge, and texture. We trained the program to distinguish between background (dark) and the droplets using manual annotations/label. For the thresholding, we used the default smoothing value (1.0 and 1.0) with 0.70 threshold. For the size filter, we put value correspond to the settings in ImageJ, 22500 for minimum size and 62500 for maximum size. This was followed with standard object selection feature option and selecting the detected droplets in object classification as a sample. After finishing the setup, we obtained the results by exporting both object predictions and measured features.

In QuPath, we started the workflow by uploading the image. Once the selected image was ready, we performed annotations similar to Ilastik. This process aimed to distinguish the background and foreground (droplets). After annotating the image, we performed “Pixel Classification” using Artificial Neural network (ANN_MLP) classifier with High (downsample = 4.0) resolution. For the features, the scales were 1.0, 2.0, 4.0 for Gaussian, Gradient magnitude, Hessian determinant and Hessian max eigenvalue. We created the object detection for droplets and measured all detected droplets. We set thick boundary class to make borders between each of the droplets. We saved the measurement data from measurement menu.

### Batch processing from each of the software

In CellProfiler, we performed batch processing by loading set of images in the Images module and run the “Analyze Images” button. For ImageJ, we executed batch processing using “Batch Process” option under “Process” tab. We used recorded macro with some adjustments to execute the images in Input folder. By processing the images through this option, we generated results directly to the Output folder. In Ilastik, we continued the batch processing straight after setting up the workflow. Similar to CellProfiler, we executed the batch processing after uploading the images and only needed to press “Process all images” button. In QuPath, we transformed the workflow from single image into scripts to execute the batch processing. Since QuPath provides the script builder, we did not have to script by ourselves and we could start batch processing by executing the script and ran it for the whole image set in the project. However, the image results from QuPath require additional script using Groovy. We managed to generate the results and you may find the script in our Github. We store both single and batch processing pipeline from each of the software here: (github.com/taltechmicrofluidics/Software-Analysis).

### Data acquisition and processing

We gathered all of the results and processed them in Microsoft Excel as follows. We tested the results with sensitivity and specificity test and used manual counting as a golden standard^36,65,66^. We used these formulas for the test:

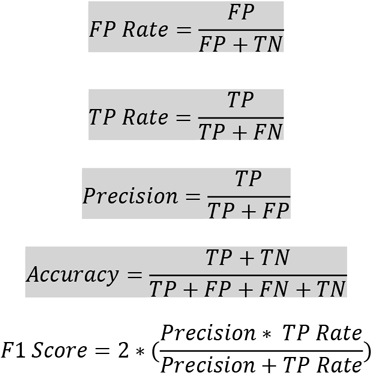

TP = Correct droplet detection compared to ground truth
FP = Wrong detection (detecting background)
FN = Wrong detection (software cannot recognize existed droplet)
TN = Background (0)
Accuracy = Quality of correctness
Precision = Similarity upon repeatable counting
Fl Score = Test accuracy measurement in dataset

## AUTHOR INFORMATION

### Author Contributions

The manuscript was written through contribution of all the authors. IS, OS and OPS conceived the study. IS conducted the research and wrote the article with support from all the other authors. SB and PP were responsible for the microbiology part, droplet experiments and microscopy imaging. All authors have given approval to the final version of the manuscript.

## ACKNOWLEDGMENT

The research was performed partially in the laboratories set up with the support from TTÜ development program 2016-2022 (project 2014-2020.4.01.16.0032). We also acknowledge the Estonian Research Council grants MOBTP109, PRG620 and MOBJD556. Authors received help for early software screening from Matin Nuhamunada, Afif Pranaya Jati and Ahmad Ardi in Universitas Gadjah Mada (Indonesia).

