## Supplementary material for "Investigation of Different Free Image Analysis Software for High-Throughput Droplet Detection": Supp.

### ASSOCIATED CONTENT

#### Supporting Information

**Supplementary Table S1. Diagnostic test on droplets detection.** We counted True Positive (TP), False Positive (FP), False Negative (FN), and True Negative (TN) by comparing the detected droplets (7145) with manual counting. For the accuracy, precision, FP rate, TP rate and F1 score, the methods are described in the manuscript.

| Output | ImageJ | CellProfiler | Ilastik | QuPath |
|---|---|---|---|---|
| Accuracy | 92.72% | 96.23% | 74.70% | 80.92% |
| Precision | 96.27% | 99.80% | 80.21% | 83.14% |
| TP | 6892 | 6885 | 6338 | 7032 |
| FP | 267 | 14 | 1564 | 1426 |
| FN | 274 | 256 | 583 | 232 |
| TN | 0 | 0 | 0 | 0 |
| FP Rate | 97.45% | 5.47% | 268.27% | 614.66% |
| TP Rate | 96.34% | 93.72% | 85.27% | 85.27% |
| F1 Score | 0.98 | 0.97 | 0.88 | 0.88 |

**Supplementary Table S2. Hardware requirement from each software.** Each of the software has its recommended requirement to make the program run smoothly during the image analysis.

| Requirement | CellProfiler | ImageJ | Ilastik | QuPath |
|---|---|---|---|---|
| Version | 4.0.3 | 1.52p | 1.3.3 | 0.2.3 |
| Operating System | Win, Mac | Win, Mac | Win, Lin, Mac | Win, Lin, Mac |
| Bit Machine | 64 and 32 | 64 and 32 | 64 | 64 |
| RAM | 4 Gb | NA | 8 Gb | 4 Gb |
| Hard disk space | NA | NA | NA | 500 Mb |
| Programming language | Python | Java | Python | Python |
| More | - | OpenGL 1.3<br>MacOS X 10.4<br>Not supported 32bit MacOS | For 3D sample, 32 Gb is expected for smooth interaction | MacOS requires at leastOS X 10.7.4 |

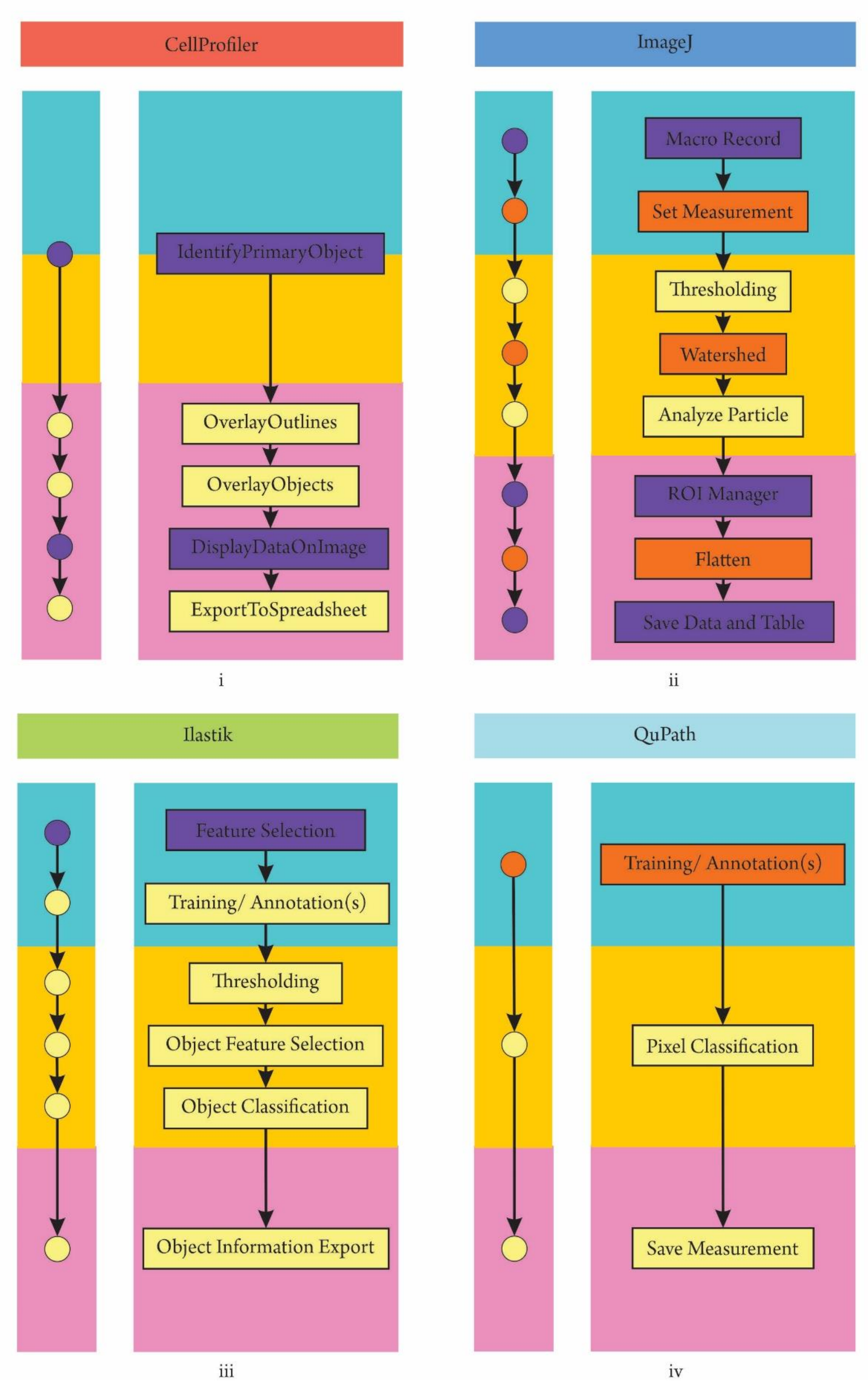

**Supplementary Figure S1. Brief workflow/pipeline from third level of complexity from most popular software** i) CellProfiler, ii) ImageJ, iii) Ilastik, and iv) QuPath.

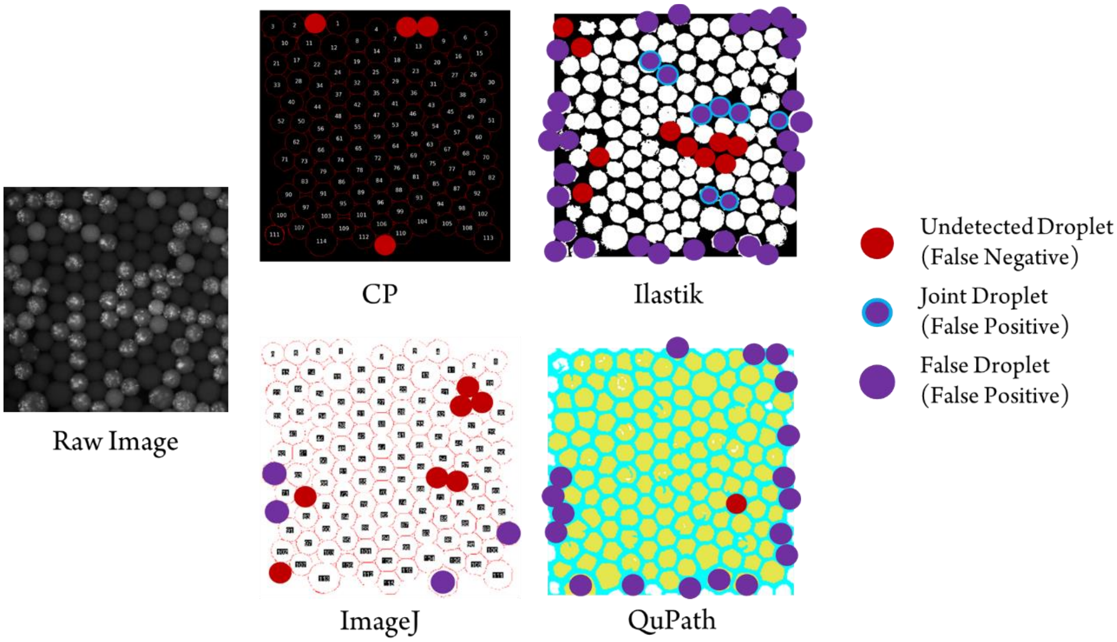

**Supplementary Figure S2. Diagnostic test in CP (CellProfiler), ImageJ, Ilastik, and QuPath.** The wrong or false detection of droplet is marked using the red and purple dots. All false detection where software unable to detect droplet is classified as False Negative/FN and colored as red dot. For the wrong detection where actually droplet is not meant to be counted, i.e. touching border, joint droplet, and wrong detected droplet, will be classified as False Positive/ FP.

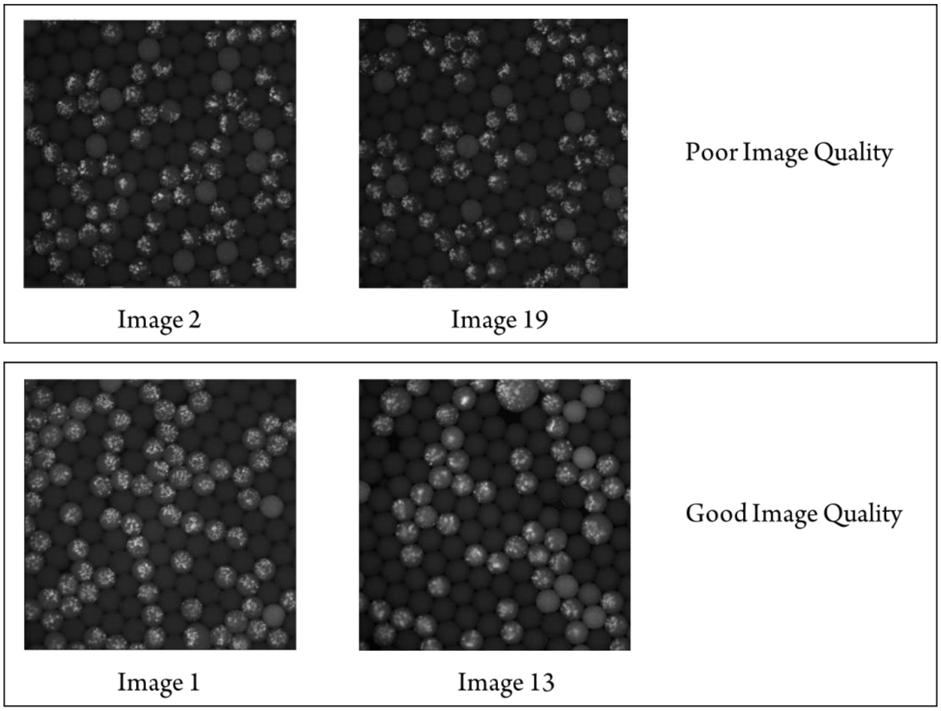

**Supplementary Figure S2. Different image quality in the experiment.** The errors were higher in poor image quality, which cannot be distinguished easily with bare eyes.

#### Step by step in CellProfiler (CP)

##### Starting

1. We used our previous pipeline of CP as the basis of this pipeline construction.
2. Referring to our previous article, you could find the full guideline, including the starting-up and explanation of CP in the supplementary here: https://doi.org/10.1039/D0AY00031K
3. Here, we only simplify the pipeline and add more modules.

##### Making pipeline in CP

4. Make sure you have opened CP and see the main GUI.
5. Once the CP is ready, you can start to set up some modules.
6. It is a default that you will have four standard modules in the pipeline (**Images**, **Metadata**, **NameAndTypes**, and **Groups**) and you can find the detailed explanation in our previous article.

##### Upload images in Image module

7. First, you need to upload your images in **Image** module.
8. For this experiment, in the **Image** module, we uploaded the 64 images of droplets which obtained from fluorescent microscope.
9. For uploading the images, you can double click the "Drop files and folders here" and find your images or just drag and drop your folder/ images there.
10. Make sure you can find the images in the box and try to see it by double click the name file.
11. By using right click on the files, you can play around, e.g. measure distances, show image histogram, see different image contrast, do interpolation, and save subplot.

##### Setup metadata module

12. This setup will help you to set the data structure in CP. For the setting, we used this following setting:
    a. Extract metadata? -> "Yes"
    b. Metadata extraction method -> "Extract from file/ folder names"
    c. Metadata source -> "File name"
    d. Regular expression to extract from file name -> "Slide(?P<Site>[0-9][0-9]_"
    (We have our images the same name format from our previous article).
    e. Extract metadata from -> "All images"
    f. Metadata type -> "text"

##### NamesAndType module

13. This module helps classifying data type into some names which may be used later in the software. We used these following settings to name our objects of interest (Bacteria):
    a. Assign a name to -> "Images matching rules"
    b. Proceed as 3D -> "No"
    c. Match -> "All" of the following rules
    d. The criteria follow "File"-> "Does"-> "Contain"-> "Bacteria" (***Critical note***: remember for not having space for the criteria name)
    e. Name to assign these images -> "FluorescenceImage"
    f. Select the image type -> "Greyscale image"
    g. Set intensity range -> "Image metadata"

**Groups module**

14. We did not use any grouping in this pipeline.
    a. Do you want group your images -> "No"

**IdentifyPrimaryObject module**

15. We also used similar setting for this module:
    a. Use advance settings -> "Yes"
    b. Select the input image -> "FluorescenceImage"
    c. Name the primary objects to be identified "Droplets"
    d. Typical diameter of objects, in pixel units (Min, Max) -> "150 – 250"
    e. Discard objects outside the diameter range? -> "Yes"
    f. Discard objects touching the border of the image? -> "Yes"
    g. Thresholding strategy -> "Global"
    h. Manual threshold -> "0.023"
    i. Thresholding smoothing scale -> "0" (in CP 3.1.8, this setting disappear once we selected manual threshold. In CP 4.0.4, this option is available, and we select "0" to make no smoothing, similar to our previous pipeline).
    j. Method to distinguish clumped object -> "Shape"
    k. Method to draw dividing lines between clumped objects -> "Shape"
    l. Automatically calculate size of smoothing filter for declumping -> "No" with "0" size of smoothing filter. (In previous software version, this option was not available. Therefore, we are not using this option in for this pipeline).
    m. Automatically calculate minimum allowed distance between local maxima -> "Yes"
    n. Speed up by using lower-resolution image to find local maxima -> "Yes"
    o. Display accepted local maxima -> "No" (The previous CP did not have this option as well).
    p. Fill holes in identified objects? -> "After decupling only"
    q. Handling of objects if excessive number of objects identified -> "Continue"

**MeasureObjectSizeShape module**

16. We used this module to measure the droplets' size and shape. In our previous pipeline, we did use MeasureObjectIntensity because we needed the pixels intensity database for making classifier in CPA. In this pipeline, we aimed to use CP as a standalone image processing software in comparison to other available software. For the setting, we used the following options:
    a. Select object sets to measure -> tick "Droplets" (Since we only detect droplets, this will only show droplets. If you have more objects to detect, you can choose any object to measure).
    b. Calculate the Zernike features -> "No"
    c. Calculate the advanced features -> "No"

**ExportToSpreadsheet module**

17. We used this module to obtain the measurement data. We implemented this following settings:
    a. Select the column delimiter -> Comma (",")
    b. Output file location -> "Default Output Folder" (or folder you prefer to save the work)
    c. Add prefixes to files names? -> "No"
    d. Overwrite existing files without warning -> "No"
    e. Add image metadata columns to your object data file? -> "No"

f. Add image file and folder names to your object data file -> "No"
g. Representation of Nan/ Inf -> "NaN"
h. Select the measurement to export -> "Yes"
i. Press button to select measurement (click the button and select "All")
j. Calculate the per-image mean values for object measurements -> "No"
k. Calculate the per-image median values for object measurements? -> "No"
l. Calculate the per-image standard deviation values for object measurements? -> "No"
m. Create a GenePattern GCT file? -> "No"
n. Export all measurement types? -> "Yes"

***Critical note***: Once the measurements were completed, we added three more modules to export the image. This image will contain the number of detected droplet based on CP's counting. We used this result to perform the diagnostic test for CP compared to the ground truth (manual counting).

**DisplayDataOnImage module**

18. This module will help you to count each of the detected droplet. The settings were:
    a. Display object of image measurements? -> "Object"
    b. Select the input objects -> "Droplets"
    c. Measurement to display -> ("Category" -> "Number") and ("Measurement" -> "Object_Number")
    d. Display background image? -> "No" (If you want to display the background, you can select "Yes")
    e. Select the image on which to display the measurements -> "FluorescenceDroplets"
    f. Display mode -> "Text" (it also has option as color, similar to the results in "IdentifyPrimaryObject"). For our case, we selected "Text" for showing the numbers in each of the droplets. We performed this to conduct diagnostic test.
    g. Font size (points) -> "32" (this can be adjusted into any number depends on the preferences.
    h. Number of decimals -> "0"
    i. Annotation offset (in pixels) -> "0"
    j. Name the output image that has the measurements displayed -> "Number" (***Critical step***: This name will be used for the next module "OverlayOutline". Therefore, do NOT mix the name with any name in the previous modules. Otherwise, the module will not have correct image data).
    k. Image elements to save -> "Image" (if you need a figure, this setting may allow "Figure" as an option).

**OverlayOutlines module**

19. We used this module to add the outline of each of the droplets with numbers we have set previously. The settings were:
    a. Display outlines on a blank image? -> "No"
    b. Select image on which to display outlines -> "Number"
    c. Name the output image -> "Outline" (***Critical note***: This name will be used in "SaveImages". Therefore, remember to give the name that represents the module).
    d. Outline display mode -> "Color"

e. How to outline -> “Thick” (There are two more option which are “Inner” and “Outer”. However, we prefer to have this option to show better outline).
f. Select objects to display -> “Droplets”
g. Select outline color -> “ ” (you may choose your preferred color for the outline by clicking the color box).

**SaveImages module**

20. Once the figure met our expectation, now we could save the image. For the settings, we used:
    a. Select the type of image to save -> “Image”
    b. Select the image to save -> “Overlay” (This refer the file we generate from “OverlayOutlines” module)
    c. Select method for constructing file names -> “From image filename” (You can pick the method for this option from Metadata as well).
    d. Select image name for file prefix -> “FluorescenceDroplets
    e. Append a suffix to the image file name? -> “No”
    f. Saved file format -> “tiff”
    g. Image bit depth -> “8-bit integer”
    h. Output file location -> “Default Output Folder”
    i. Overwrite existing files without warning? -> “No”
    j. When to save -> “Every cycle” (***Critical note***: This setting will not dependent on the previous cycle in the pipeline. If you want to make an aggregate image, there is “First cycle” and “Last cycle” option).
    k. Record the file and path information to the saved image? -> “No”
    l. Create subfolders in the output folder -> “No”

**Testing and Batch Processing**

21. To test your module, click “Start Test Mode” and “Run”. You can also run the modules step by step using “Step” button. The “Next Image Set” is option if you want to run the pipeline for another image which you have prepared in **Images** module. If you finish testing the pipeline, click “Exit Test Mode” and you can start analyzing the whole set of images.
22. To run the batch processing or analysis, simply click “Analyze Images”.
23. The full CP pipeline for this project can be found here:
    https://github.com/taltechmicrofluidics/Software-Analysis/CP/

#### Step by step in ImageJ

##### Starting-up the software

1. Download the ImageJ software and you can find the software here: [https://imagej.nih.gov/ij/download.html](https://imagej.nih.gov/ij/download.html)
2. Install the software in your computer by double clicking the ImageJ installer you have downloaded through the link.
3. Open ImageJ software (double click the icon or right click and select open).
4. You will find the interface below once the program is starting.

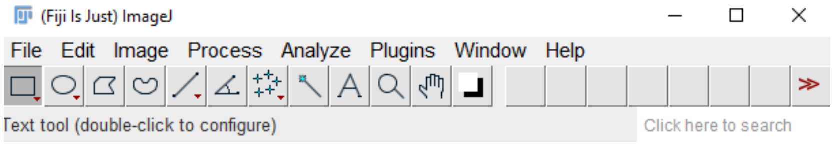

##### Pipeline in ImageJ

5. Once the program is ready, you can start using ImageJ for detecting droplets.
6. You can make the ImageJ's window as "full screen" to make the window more visible.

##### Upload Image

7. You can start uploading the image by dragging your image file to ImageJ. To upload the image, you can also click "File" and select "Open ... " and the program may allow you to select image from the software.
8. ***Critical step***: Once you have your image opened, find "Macro record" through Plugins -> Macros -> Record. This feature of the software will allow you to record any activity in ImageJ and will be useful for running the batch processing later.
9. ***Critical step***: If you only want to proceed with one image, "Macro record" is not mandatory. If you need to perform batch processing later, we recommend you use this option to ease the macro script later. Duplicate your image using "Image" and select "Duplicate ... " or using simple keyboard command Ctrl+Shift+D and give name for your duplicate images. For our case, we used two duplication. First, the duplication will be used to construct the outline of measurement and the second duplication will be used to measure the pixels intensity. For the first, we named the image as "droplets" and the second one we named it as "measure".
10. To make sure your working tab is the image you prefer, click the image once again. Here, we select the tab of "droplets" as our main working image. ***Critical note***: the working tab will receive the command from the next option you pick from the software. As an example, if you have multiple images to work with (see below), your working tab is the one with black title.

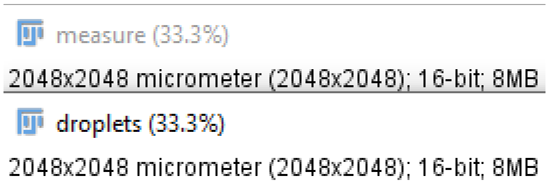

##### Thresholding option

11. Once you have your images ready, you can give commands to implement thresholding in your image.
12. Click Image -> Adjust -> Threshold ... or you can just simply press Ctrl+Shift+T on your image.

13. Threshold window will show up, and you can adjust your threshold there. For our case, we tried to implement similar thresholding scale 0.023 in CP. Therefore, we used "Default" and simple calculation 0.023 x 65535 = 1507 as minimum the threshold. We had 16-bit image, therefore, the 65535 represents the total pixel value within an image. If you have 8-bit image, the scale would be ranging from 0 to 256.

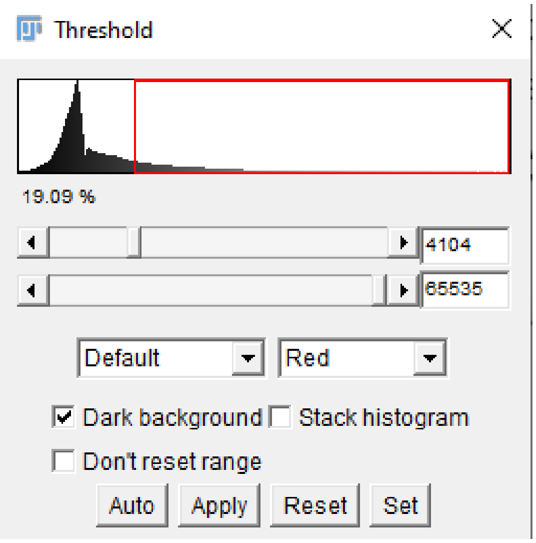

14. The Threshold tab also provide options to simplify the thresholding method, such as Huang, Otsu and etc.
15. The thresholding visualization also provide three options, such as Red, B&W, and Over/Under.
16. ***Critical note***: Do NOT forget to tick "Dark background" if you have bright background, untick this option.
17. If you have multiple images and want to implement the thresholding at the same time, tick the "Stack Histogram". ***Critical note***: If you want to run batch processing later, it is better to find one thresholding which work best for most of the image and implement the same scale. This stack histogram thresholding would make the analysis harder later.
18. Keep the "Don't reset range" untick if you did not do any brightness and contrast adjustment. This setting will reset the range. Unless, you prefer to keep the threshold range after some adjustment in brightness and contrast adjustment.

**Watershed option**

19. Since the pixels between droplets are vaguely distinguishable, the classification would simply put the pixels as foreground. As the results, the droplets look connected to each other. This Watershed option is used to perform the segmentation based on this site.
20. The option is simple, once you have the thresholded image, find in tab Process -> Binary -> Watershed
21. Once you click the Watershed, you will find each of the droplets.

**Analyze particle option**

22. Before executing this option, you need to set the measurement template image in the "Analyze Particle" option. This can be done using tab "Analyze" -> "Set Measurements … ". Here, we ticked Area, Standard deviation, Mean gray value, Centroid, Perimeter, Median and Display Label. For the "Decimal places", we used the default number "1". ***Critical step***: Select the "Redirect to" into your image for measuring the pixels intensity, in our case, we have named the image as "measure".

23. You now have ready image for the analysis. You can directly perform the droplet detection and counting through Analyze -> Analyze Particles …
24. When you click the option, another tab will appear (below).

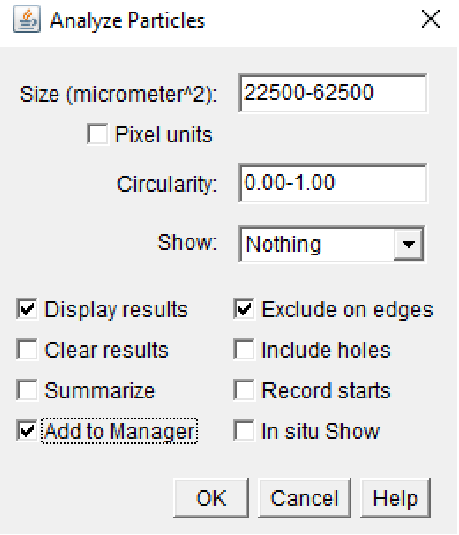

The Analyze Particles option provides some boxes to set the preferred size, circularity, and etc.

a. For our droplet detection, we use the size limit we used in CP. We limit the 150-250 μm for the diameter of droplets in CP's pipeline. Here, we put the number as the square large of object (droplet) in $\mu m^2$. Therefore, the limit is 22500-62500 $\mu m^2$
b. We put 0 – 1 circularity. Note: Only ImageJ which require the circularity and we put the range as the default from the program.
c. We select "Outline" for showing the result.
d. For showing the results, tick "Display results"
e. The "Exclude on edge" option is to make sure we do not include droplets which touch the border of the image.
f. The "Add to Manager" will add the counting in the tab which appear after the analysis occur ("ROI Manager").
g. We did not use any other option since they were not needed for our analysis.

**Managing the results**

25. You will find your results (Droplets in outline and a table) with one tab of "ROI Manager". This tab is used to maintain the result.
26. For obtaining the droplet image, click "Flatten". Once this button is pressed, you will have another image window. ***Critical note***: before acquiring the image, find "More" and set the number's font in "Labels … " into any scale you may prefer. Click "File" in the main menu and save the image using "Save As" with your preferred format, in our case, we picked .TIFF file format.
27. For the table, you can directly save the table from its menu. Using "File" and "Save As … " and type the title with .csv ending. Critical note: The table will be saved as .txt file if you do not put any specific filetype. The .csv can be used in MS Excel or any numerical analysis software. The example of results in ImageJ can be seen below.

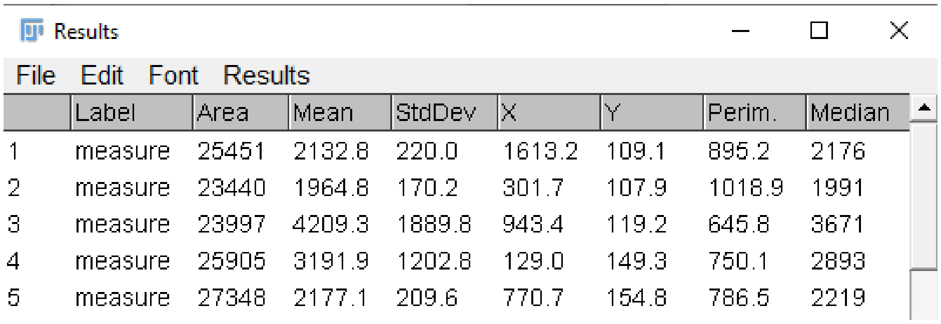

| | Label | Area | Mean | StdDev | X | Y | Perim. | Median |
|---|---|---|---|---|---|---|---|---|
| 1 | measure | 25451 | 2132.8 | 220.0 | 1613.2 | 109.1 | 895.2 | 2176 |
| 2 | measure | 23440 | 1964.8 | 170.2 | 301.7 | 107.9 | 1018.9 | 1991 |
| 3 | measure | 23997 | 4209.3 | 1889.8 | 943.4 | 119.2 | 645.8 | 3671 |
| 4 | measure | 25905 | 3191.9 | 1202.8 | 129.0 | 149.3 | 750.1 | 2893 |
| 5 | measure | 27348 | 2177.1 | 209.6 | 770.7 | 154.8 | 786.5 | 2219 |

**Batch processing**

28. By using the pipeline, you may be able to detect and gather the results for single image. For batch processing, you need to set the setting using macro script.
29. Find the macro record script tab which you have opened since the beginning of the process and you will receive these lines of script (see below).
 You have everything recorded in this tab, therefore, you need to clean some unnecessary line e.g. "selectWindow("0.tiff")" which generated when you click you working tab (image) or "run("Close") that represent closing a tab.
30. The script cleaning is depending on the user. ***Critical note***: It is not a complex programming, but you still need to take a note for every punctuation, such as ; and " are important.
31. For this droplet detection, we cleaned some scripts below.

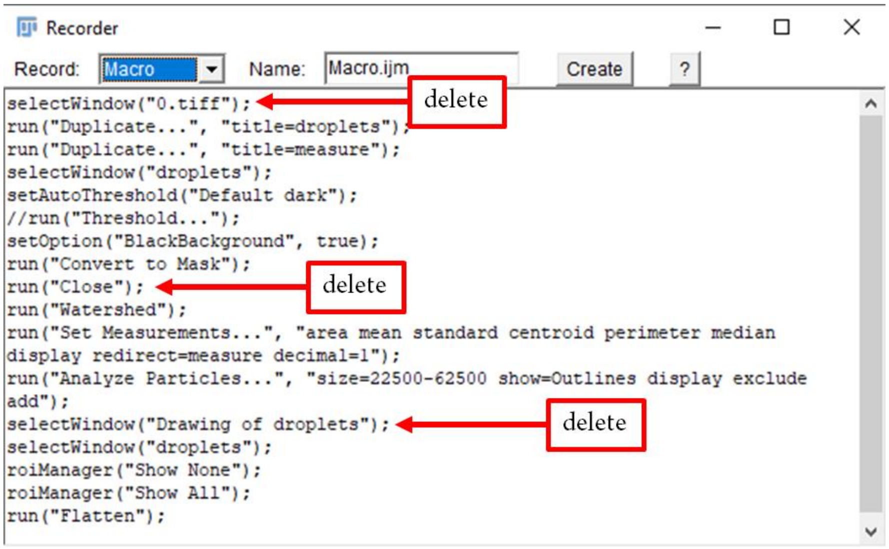

32. When you finish cleaning up the unnecessary lines, you can save your script by clicking "Create". You may assign a name for this macro. We used default name as Macro.ijm. ***Critical note***: The .ijm is mandatory for referring that the macro is for ImageJ.
33. You can find our final macro script in our Github with .ijm format here: https://github.com/taltechmicrofluidics/Software-Analysis/ImageJ/
34. Once you have the script, you can run your batch processing by opening "Process" -> "Batch" -> "Macro... "
35. You will find new tab called Batch Process. Here, you need to adjust some settings.
    a. First, you need to define your input folder. By clicking the "Input... " you can select where you have the images you want to process.

b. Second, the output folder. You can define the folder by clicking the "Output… " button/
c. For the output format or third, you can select various file type. In our case, we prefer to have "TIFF" as the output format.
d. For the "Add macro code", let it be as "[Select from list].
e. We also emptied the "File name contains"
f. You will find another empty tab below the "File name contains" and you can copy your script into this white tab. Or, if you saved the .ijm macro file, you can click "Open… " to open your macro script. If you change some lines within this tab, you can also save the script through "Save… " tab.
g. You can test your script using "Test". If a table result pops out, then the script may work as you expected. The image will also appear during the test, but not the processed images. You will find the processed image in the output folder after you click "Process".
h. The table results will accumulate the counting for the whole folder. Critical note: Do NOT close the results tab when the analysis starts. Once the analysis is finished, you can save the results through "File" and "Save As… ". You can name the results with .csv as the filetype.

#### Step by step in Ilastik

##### Starting the software

1. Find and download the Ilastik ver. 1.3.3 software here: https://www.ilastik.org/download.html
2. Install the file into your directory (follow the instruction from the software developer).
3. Once the software is installed, you will find the interface below.

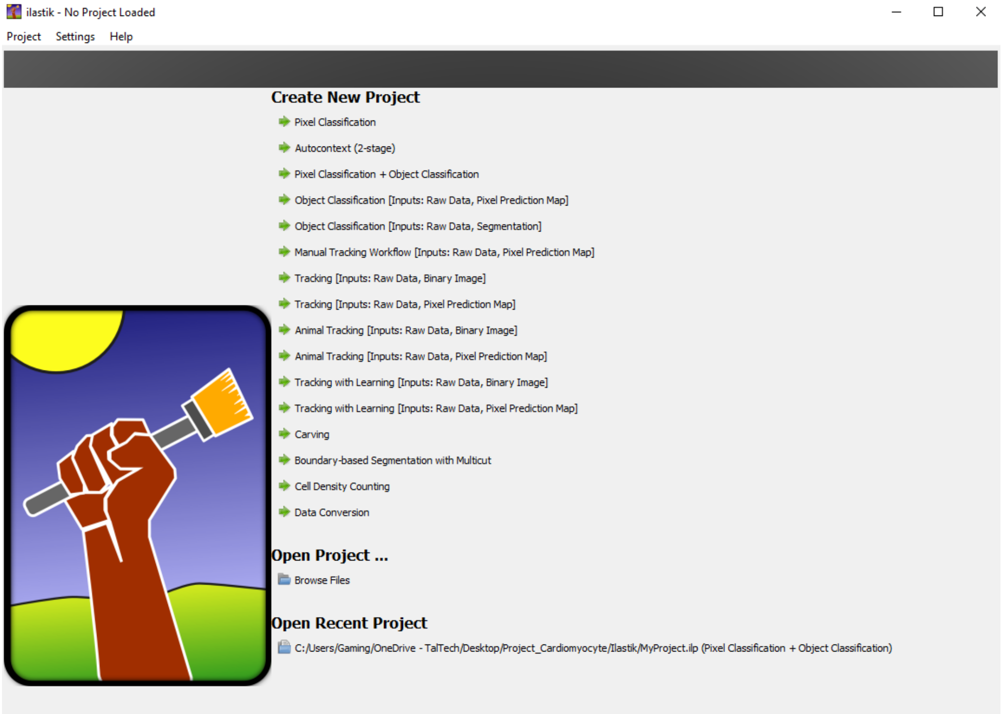

4. You can find the predefined pipeline which provided by Ilastik in the main interface. In our case, we will use the "Pixel Classification + Object Classification". ***Critical note***: before clicking the pre-defined pipeline, prepare a folder which may contain your image and a directory to save Ilastik's file. Once you set the folder, do NOT move, or change the directory to minimize error in opening your project.
5. Click the "Pixel Classification + Object Classification" and give your project's name and your directory. Once you click save, you will be redirected into Ilastik's main user interface with selected predefined blocks.
6. In "Pixel Classification + Object Classification", you will have nine blocks on the left side which represent each module for each steps.

##### Data Input module

7. For uploading the image in your workspace, you can find the 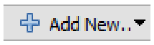

in the "Raw Data" tab on the right sight of the tab. If you have "Mask" for your data, you can also add the mask in "Atlas" tab.
8. Once you select your image, you can find the image will appear on the screen along with some buttons. You can adjust the size of the figure using these buttons 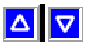 . If you need to rotate or flip, you can do it using these buttons 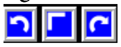 . We only adjusted the size for seeing the image conveniently.
9. Once the image is uploaded, you can proceed to the next module, "Feature Selection", by clicking the module.

**Feature Selection module**

10. For the feature selection, there are some options which you can add to help the training later. By clicking the "Select Features", you will find three adjustable scale for Color/ Intensity (using Gaussian Smoothing), for Edge (Laplacian of Gaussian, Gaussian Gradient Magnitude, and Difference of Gaussian), and for Texture (Structure Tensor Eigenvalues and Hessian of Gaussian Eigenvalues). These features selection may give different effect during the training later, such as edge enhancement or blur. However, we received the suggestion from www.image.sc to use three scales for each of the setting, which are 0.3, 1.00, and 3.50. You can select these simply by clicking the box under the number. The software also provide the empty scale if you want to optimize the features by adding on the "add" box next to 10.00 or change any available number. Make sure you tick to choose the scale.

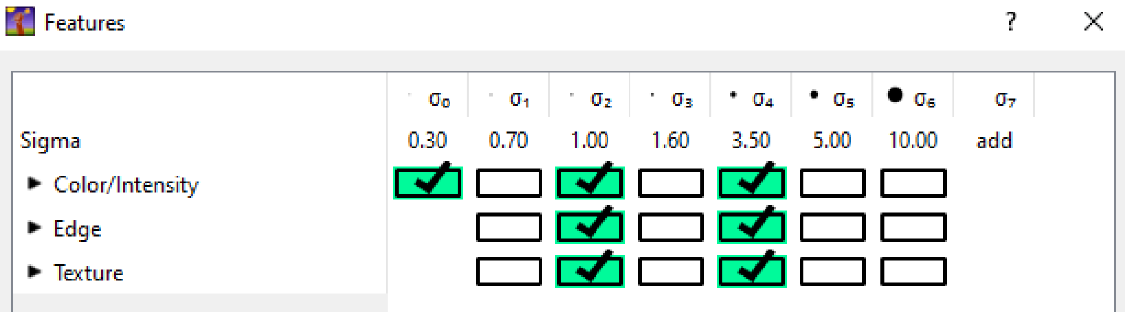

11. After selecting the features, you can proceed to "Training" module.

**Training module**

12. In the "Training" module you need to label your objects. Here, we define two types of object. First, we want to have droplets and the second we define the background. In other software, this feature is performed by making a thresholding and distinguish between the foreground (droplets) and background (dark). In Ilastik, you need to label or annotate the object with paints. You can add label by clicking or use available Yellow and Blue. 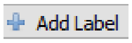

You can change the name by clicking the name next to the color box. You can also change the color you may prefer by double clicking the color box.
13. Once you finish setting up the labels name and color, you can start annotating your image. We use Yellow to annotate the droplets and Blue for the background (see below).

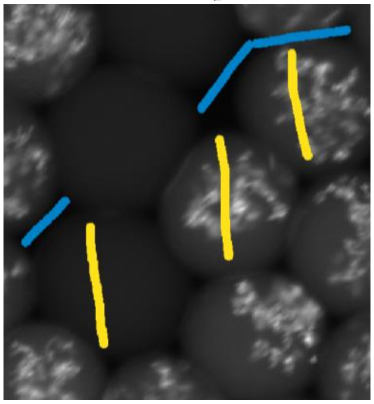

14. If by mistaken you put the wrong annotation, you can erase the label using this button and re-annotate the label you want to remove. It is like making an overlay but this will erase your annotation. 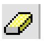
15. You can check if your annotations work best for the image by clicking "Live Update" and directly see the changes. You can also do more annotations when putting the processing in this option.
16. If you are satisfied with the training, you can proceed to the "Thresholding".

**Thresholding module**

17. This module provide smoothing tool, thresholding option and size filter. For this module, we use these following setting:
    a. Method -> Simple
    b. Input -> Yellow box
    c. Smooth -> 10.0 and 10.0
    d. Threshold -> 0.60
    e. Size filter -> Min (11250) and Max 62500
18. After putting the setting, click "Apply" for implementing the setting and you will see the example of the result (below).

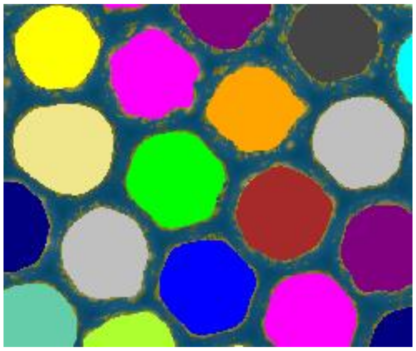

19. Now you will proceed to the next module, "Object Feature Selection".

**Object Feature Selection module**

20. In this module, you need to pick the features using "Select Features" similar to the previous one. In this case, this feature will extract the data from the object. Critical note: when selecting the features, we recommend using the box on the left bottom "All excl. Location" and press OK. This feature selection will gather the results from each of the detected object, e.g. intensity distribution, shape size, and etc.

**Object Classification module**

21. For this module, you just need to select the droplets using one of the label (similar to the "Training" module).
22. We do not need to distinguish between one droplet and another. Therefore, here, you just need to select the droplets with one label (see below).

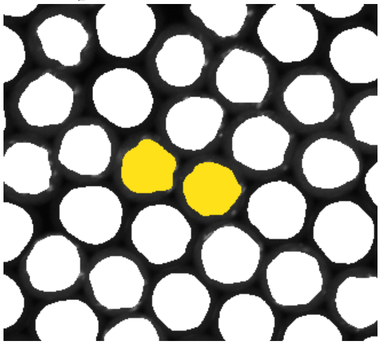

23. This will allow the software to take account all of the droplets when exporting the data.

**Object Information Export**

24. This module allows the results into figures and tables.
25. To export the image, you need to set the "Source" as "Object Predictions".

26. For exporting the table result, click the "Configure Feature Table Export". We set our table to be in .CSV format. You can change the format by selecting the .CSV in the drop-down selection in General.
27. At the bottom, you can find the box "Features" to select the data you prefer to have. To gather all of the data, click "All" and "OK"
28. Once you finish with the setting, just hit the "Export All" button and you will find the results in your folder.

**Batch Processing**

29. For running a batch analysis, you just need to click "Batch Processing" and upload all of your images.
30. Once you upload the images using the "Select Raw Data Files" button, you can press "Process all files" to run the batch processing.
31. The Ilastik pipeline can also be downloaded here: https://github.com/taltechmicrofluidics/Software-Analysis/Ilastik/

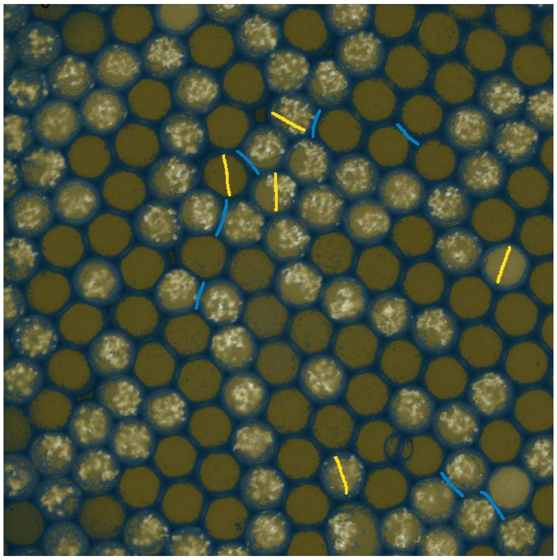

#### Step by step in QuPath

##### Starting the software

1. Download the QuPath software version 0.2.3 from https://qupath.github.io/
2. Follow the instruction and guideline for installing the software
3. Once you finish the installation, open QuPath from your desktop
4. When opening QuPath for the first time, the software will ask the memory allocation. For this case, you can choose 4 Gb or less. Make sure you have enough memory to run your Windows and other program. For the Numbers and Date, we use the default English (United States). If you prefer, you can tick the check update as well. Click Apply to implement the setting.
5. Now, you should have the QuPath main interface.
6. You will have the main tab and tools at the top (see below). The main tab contain necessary tools which will be used later, especially the "Measure" and "Classify". The tools are for annotating the object of interest.

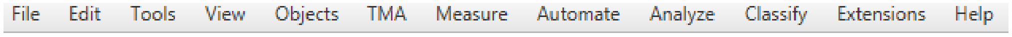

##### Making project and upload images

7. Now, we are focusing on the use of QuPath for droplet detection. For starting the project, select File and Project to create new project. ***Critical note***: Prepare a folder with your raw image in it. Do NOT move any content from this folder to minimize any error later.
8. Once you create project, you will have the working window (below).

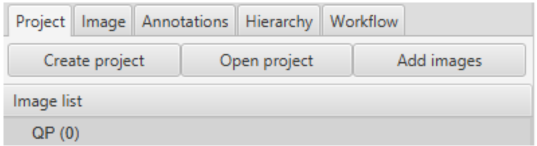

9. To upload your images, click "Add images" and select your images. ***Critical note***: We recommend selecting all of your images and upload all of them now. Even though you will be annotating and classifying the droplets in one image, this will make the workflow easier later. When selecting the image, you can drag and drop the file on the box or there are other choices, such as "Choose files" from the folder, "Input URL" if your files are available online, "From clipboard" if you are copying through your operating system, or "From path list" if you are calling the image from the whole folder. Once you have the images, you need to set these following settings:
   a. Image provider -> "Default (let QuPath decide)
   b. Set image type -> "Fluorescence"
   c. Rotate image -> "No rotation"
   d. Press Import to put the image in your project
10. You will find all of the image in the Image list and you are ready to annotate the image.

##### Image Annotation

11. This image annotation will help the pixel classification into desired object of interest (in our case droplets).
12. Select one image and double click the image. You may find the image on the right side of the list.

13. Now click the "Annotations" tab in working window and the (see below).

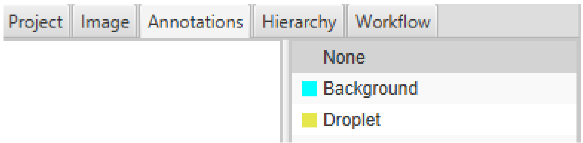

14. Before annotating the image, you need to add classes. In our case, we aim to classify the pixels into "Background" and "Droplet". You can add the class using " : " button next to "Auto set" or just press *right click* on the class box. Click "Add/Remove" and select "Add class". You can have any name and press "OK". In our case, we have two classes: Background (Cyan) and Droplet (Yellow). If you need to delete the class, you can click the class and perform the right click and select the "Remove class" from "Add/Remove".
15. Once you have the class, you can change any color you may prefer for your class by clicking the color box next to the class name.
16. For the annotation, you can use these buttons 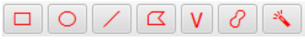 to select any pixels and assign the annotation into specific class.
17. After making the annotations, set each of the annotation into the class you may want to be using "Set class", e.g. the circles as Droplet and the irregular form as Background. You will see if the annotations are correct, you will have the number after the class (example below).

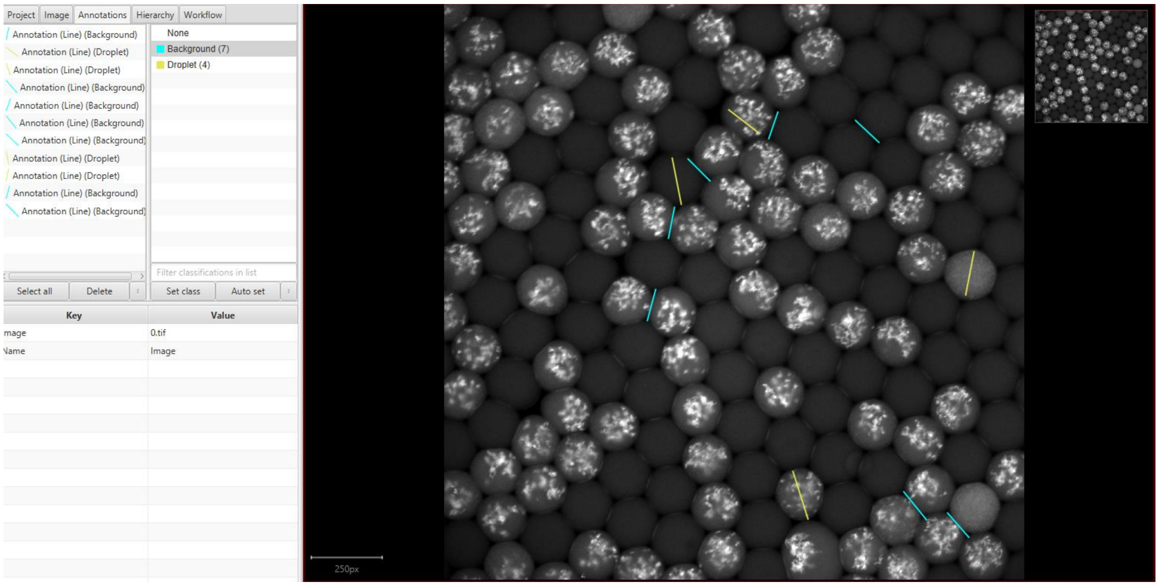

18. When you finish annotating the image, you can proceed to pixels classification step.

**Pixel classification and measurement**

19. You can find the pixel classification option under "Classify" in main tab. Select Pixel classification and "Train pixel classifier" or simply press Ctrl+Shift+P on your keyboard. In this option you may follow this settings:
    a. Classifier -> "Artificial neural network (ANN_MLP)
    b. Resolution -> "High (downsample = 4.00)
    c. Features -> "Default multiscale features". ***Critical note***: click "Edit" and change the "Scales" into 0.5, 1.0, and 4.0 and in "Features" select "Gaussian", "Laplacian of Gaussian", "Gradient magnitude", "Structure tensor max eigenvalue", "Hessian Determinant", "Hessian max eigenvalue". Click "OK" after selecting these.

d. Output -> "Classification"

e. Region -> "Everywhere"

f. Click "Advanced options" and set the "Boundary strategy" for having "Boundary strategy: Classify as Background" and "Boundary thickness" with "10" pixels.

g. You can see how the classification run by clicking the "Live prediction".

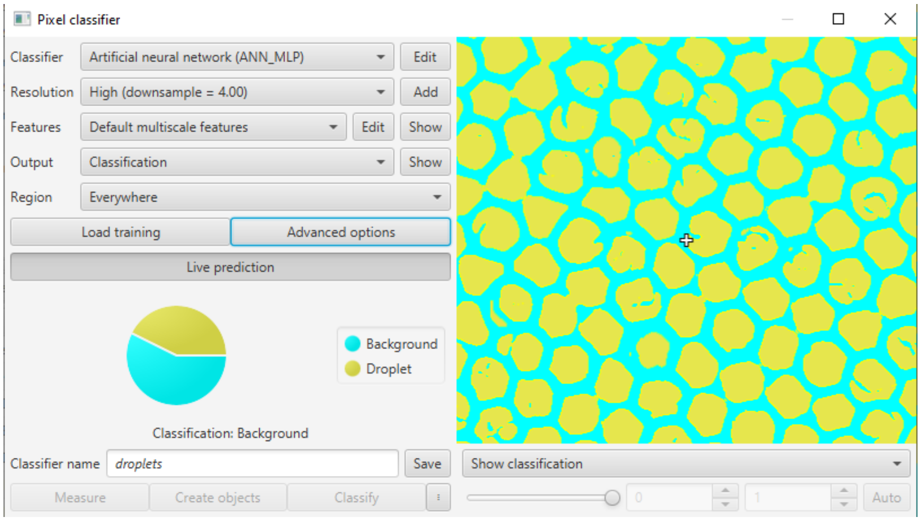

h. Name the classifier by putting the name on "Classifier name" and click "Save".

i. After saving the classifier, you need to make the pixel classification as an object using "Create objects". In this option, you need to change the "Parent objects" as "Full image" and click "OK". There will be another tab called "Create objects" which appear. For this tab, the following settings are:

   i. New object type -> "Detection"
   ii. Minimum object size -> "112500" px^2
   iii. Minimum hole size -> "0" px^2
   iv. Tick the "Split objects"
   v. Let other setting as a default and click "OK"

j. You will find that the annotations have more objects as "Droplet" followed with total number of the class.

k. Once you have all of the objects, click "Measure" button to have the counting.

l. There will be a tab appear and you just need to put "All detections" in "Select objects".

m. The measurement will start once you press "OK" button.

n. ***Critical step***: You need to perform "Create Annotations" to obtain the figure in each of the detections. If you do NOT need any image in batch processing, this step is not mandatory. To perform this, you can simply click "Classify" -> "Pixel Classification" -> "Load Pixel Classification". You already have your model of classifier "droplets". Select the model and select "Region" -> "Everywhere" and click "Create object". There will be another tab appear "Full image" and click "OK".

o. To check your measurement, you can find your measurement under main tab "Measure" -> "Show annotation measurements". ***Critical note***: You may get all of the measurement including the first annotation measurement. To filter this, you can distinguish the ROI of from your first measurement and exclude these measurements.

p. Save the measurement by clicking "Save" box and give the name for the measurement, e.g. droplet_image_0
q. For saving the image file with the label, you can simply click "File" -> "Export images ... " -> "Rendered RGB (with overlays)" and give the name for the image and click "Save".
r. Now you will have the detection, measurement, and figure for your project as single file.

**Batch processing**

20. The batch analysis requires some script writing in QuPath. Luckily, this software provide script generator (similar to macro record in ImageJ).
21. ***Critical note***: If you follow the instruction from the beginning without using other option or features of QuPath, you do NOT need any adjustment. You only need to use this following steps:
    a. Click "Workflow" tab and find "Create script" and click the button.
    b. There will be a "Script Editor" tab with some lines of script. (To confirm your script, you can find our lines of script below).

```
setImageType('FLUORESCENCE');
classifyDetectionsByCentroid("droplets")
resetSelection();
createDetectionsFromPixelClassifier("droplets", 11250.0, 0.0, "SPLIT")
createAnnotationsFromPixelClassifier("droplets", 11250.0, 0.0, "SPLIT")
selectDetections();
addPixelClassifierMeasurements("droplets", "droplets")
saveDetectionMeasurements('/C:/Users/Gaming/OneDrive - TalTech/Desktop/Project Software Review/Update Project/Article 1/Results/QP for diagnostic/')
```

    c. For running the script for whole project, you can use "Run" tab and select "Run for project"
    d. Select all of the available data and press » button and press "OK".
    e. You will find all of the data in your directory as text file. This file can be imported into MS Excel or any data analysis software.
22. ***Critical note***: QuPath can only generate the data one by one. It means, you have to check one file and another to find your detection. We used available library to combine the files and find the script written in Python here: https://github.com/taltechmicrofluidics/Software-Analysis/QuPath/Combine/
23. For obtaining the image, you have to run separate script after performing the batch analysis in QuPath. You can the script here: https://github.com/taltechmicrofluidics/Software-Analysis/QuPath/
    ***Critical step***: there are some lines which you need to consider and make sure you have correct commands.

```
def imageData = getCurrentImageData()

// Define output path (relative to project)
def outputDir = buildFilePath(PROJECT_BASE_DIR, 'export')
mkdirs(outputDir)
def name = GeneralTools.getNameWithoutExtension(imageData.getServer().getMetadata().getName())
def path = buildFilePath(outputDir, name + "-labels.png")

// Define how much to downsample during export (may be required for large images)
double downsample = 8

// Create an ImageServer where the pixels are derived from annotations
def labelServer = new LabeledImageServer.Builder(imageData)
  .backgroundLabel(0, ColorTools.WHITE) // Specify background label (usually 0 or 255)
  .downsample(downsample)    // Choose server resolution; this should match the resolution at which tiles are exported
  .addLabel('Background', 1)      // Choose output labels (the order matters!)
  .addLabel('Droplet', 2)
  .multichannelOutput(false) // If true, each label refers to the channel of a multichannel binary image (required for multiclass probability)
  .build()

// Write the image
writeImage(labelServer, path)
```
